# MAGIK: A rapid and efficient method to create lineage-specific reporters in human pluripotent stem cells

**DOI:** 10.1101/2023.09.07.556774

**Authors:** Tahir Haideri, Jirong Lin, Xiaoping Bao, Xiaojun Lance Lian

**Affiliations:** Department of Biomedical Engineering, Pennsylvania State University, University Park, PA, 16802, USA; The Huck Institutes of the Life Sciences, Pennsylvania State University, University Park, PA, 16802, USA; Department of Biology, Pennsylvania State University, University Park, PA, 16802, USA; Davidson School of Chemical Engineering, Purdue University, West Lafayette, IN 47907, USA

## Abstract

Precise insertion of a fluorescent protein into a lineage-specific gene in human pluripotent stem cells (hPSCs) presents challenges due to the low knockin efficiency and difficulties in selecting the correctly targeted cells. Here we introduce the ModRNA-based Activation for Gene Insertion and Knockin (MAGIK) approach to enhance knockin efficacy in hPSCs. MAGIK operates in two steps: first, it employs a Cas9-2A-p53DD modRNA with a mini-donor plasmid (without a drug-selection cassette) to significantly enhance efficiency; second, a dCas9 activator modRNA and a dgRNA are used to temporarily activate the successfully targeted gene, allowing for live cell sorting without single cell cloning. Consequently, MAGIK eliminates the need for drug selection cassettes or labor-intensive single cell colony screening, expediting precise genetic integration. We have demonstrated that MAGIK can be utilized to insert fluorescent proteins into various genes, including *SOX17, NKX6.1, NKX2.5* and *PDX1*, across multiple hPSC lines, showcasing its robustness. This innovative MAGIK approach streamlines the process and provides a promising solution for targeted genetic modifications in hPSCs.

## INTRODUCTION

CRISPR-mediated gene knockin methodologies have emerged as powerful tools for gene editing^1^, marking significant strides in improving efficiency, specificity, and the scope of targeting. CRISPR-Cas9 can introduce precise double-stranded breaks (DSBs) in targeted DNA sites^2–4^, thereby enabling the insertion of specific genetic variant sequences provided by the donor DNA templates. Of note, homology-directed repair (HDR) is a major mechanism that aids this process. HDR permits the precise integration of exogenous DNA sequence at a specific site in the genome where a DSB has occurred^5^. The HDR process relies heavily on donor DNA templates, which come in three primary forms: double-stranded DNA (dsDNA), single-stranded DNA (ssDNA), and adeno-associated virus (AAV). While ssDNA and AAV may enhance knockin efficiency in comparison to dsDNA, the latter is simpler to manufacture and produce.

A recent advancement in CRISPR-mediated gene knockin technology entails the integration of fluorescent proteins into lineage-specific marker genes within human pluripotent stem cells (hPSCs)^6–8^. The creation of fluorescent reporter lines has revolutionized the isolation process of desired differentiated cells and offers an invaluable tool for refining conditions to enhance stem cell differentiation^9,10^. Fluorescent reporters offer the ability to illuminate specific cell populations, thereby enabling real-time tracking of cell type specification throughout the differentiation. For instance, the knockin of EGFP into the endogenous NKX2.5 locus^11^ in hPSCs generates lines that express EGFP concurrently with the emergence of cardiac progenitors during cardiac differentiation of hPSCs. The advantage of real-time monitoring and tracking of particular cell populations lies in the opportunity it provides for uncovering the molecular mechanisms underlying the differentiation of hPSCs into defined cell types.

Due to the inherently low efficiency of knockin, a drug selection cassette (e.g., PGK-Puro^R^) was commonly included in the donor vector. This addition allows for the enrichment of correctly targeted clones. Despite this improvement, the efficiency remains relatively low, necessitating the screening of hundreds of drug-resistant clones. For example, in the case of OCT4-EGFP knockin, researchers selected 288 puromycin-resistant clones and subsequently performed PCR and Southern blot analysis to identify only 8 correctly targeted clones (2.8% efficiency)^12^. A major challenge posed by this drug-selection approach is that the drug-selection cassette interferes with the expression of the integrated fluorescent protein. Consequently, an additional step is necessary to remove the drug-selection cassette after successful knockin, which can slow down the generation of hPSC reporter lines.

To address this, the use of a drug selection cassette-free donor vector has been proposed, aiming to expedite the process. However, this selection-free approach may result in even lower efficiency. For instance, when using the OCT4-mOrange drug selection cassette-free donor vector (mini-donor)^12^, the co-transfection of a CRISPR plasmid targeting OCT4 and the mini-donor plasmid results in an efficiency of just 0.001%. This extremely low efficiency makes it impractical for working with lineage-specific genes that do not express in undifferentiated hPSCs. For lineage-specific genes, the lack of fluorescence in successfully targeted cells precludes the use of live cell FACS technology for isolating knockin cells. As a workaround, a traditional post-knockin process is adopted: cells are replated at single cell density, left for about ten days, and then single cell-derived clones are selected. A subsequent genomic PCR analysis verifies the successful knockins. However, given the low efficiency of 0.001%, researchers would be required to randomly pick over 100,000 individual colonies to establish a single accurately targeted clone, a task that is largely impractical. As a result, inserting fluorescent proteins into lineage-specific marker genes in hPSCs still remains a significant challenge.

To address these challenges, we have developed a novel strategy: <u>M</u>odRNA-based <u>A</u>ctivation for <u>G</u>ene Insertion and <u>K</u>nockin (**MAGIK**). MAGIK is designed to both enhance knockin efficiency and simplify the process of isolating successfully targeted cells. Our MAGIK method starts with the use of a Cas9-2A-p53DD modified mRNA (modRNA), an alternative to the Cas9 plasmid, to significantly enhance knockin efficiency. The subsequent step employs a dCas9 activator modRNA to temporarily activate the expression of lineage-specific gene and its associated fluorescent protein. This maneuver simplifies cell isolation through FACS, negating the need for selection of single cell-derived clones. By circumventing the necessity for the selection and genomic PCR of single cell-derived clones, our approach provides a markedly efficient and expedited alternative to conventional methods.

## RESULTS

### ModSAM system robustly activates silenced gene expression in hPSCs

CRISPR activation (CRISPRa) systems^13–15^ are extensively employed to initiate the transcription of otherwise silenced genes in a manner specific to gRNAs. In earlier work^16^, we showed how the CRISPR gene-editing efficiency in hPSCs could be increased through a modRNA-based delivery of Cas9, which resulted in up to a three-fold increase over an equivalent plasmid-based strategy. With this success, we chose to adapt our modRNA-based approach to the CRISPRa system in hPSCs. We decided to develop modRNA-based synergistic activation mediator (modSAM) system (**Fig. 1A**). Our modSAM system is comprised of two key components. The first is the dCas9 activator (VP64dCas9VP64-T2A-MPH) modRNA, which utilizes two VP64 transcriptional activators, each attached to the N and C terminus of the deactivated Cas9 (dCas9). It also includes the expression of the MS2-p65-HSF1 (MPH). The second component is an engineered ‘dead’ guide RNA (dgRNA)^17^. This dgRNA is unique as it contains two MS2 aptamers and has a spacer length of 14 base pairs (bp). A notable advantage of using this dgRNA is its ability to pair with either a dCas9 or a catalytically active Cas9 without causing DSBs. By applying our modSAM system, we managed to achieve robust activation of lineage-specific genes, NKX6.1, PDX1, and NKX2.5 in hPSCs (**Fig. 1B-D**). After modSAM transfection using PDX1-dgRNA1, 63.3% ± 3.1% PDX1+ cells were generated measured by flow cytometry at 48 hours post-transfection (**Fig 1B**). For NKX2.5 activation, we conducted qPCR on mRNA extracted from cells with modSAM transfection, demonstrating up to a 68-fold increase in NKX2.5 expression compared to the untransfected hPSCs (**Fig 1D**).

**Figure 1.**
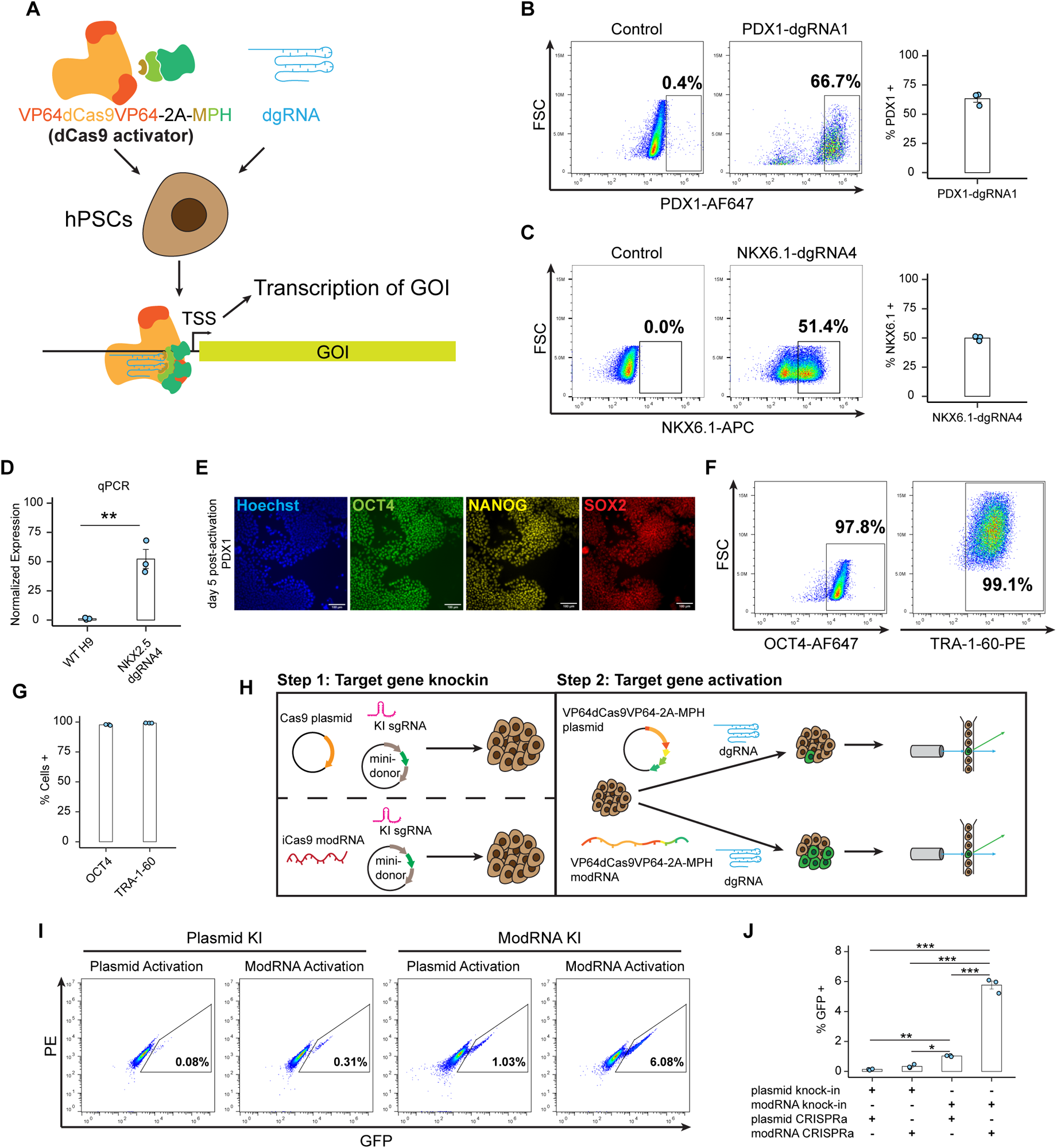
ModRNA delivery increased gene knockin and target gene activation. (A) Schematic diagram for dCas9 activator mediated activation of silenced genes in hPSCs. (B-C) WT H9 cells were transfected with dCas9 activator modRNA and either PDX1 dgRNA (B) or NKX6.1 dgRNA (C) on day 0. Cells were collected 48 hours after transfection and analyzed for PDX1 (B) or NKX6.1 (C) expression by flow cytometry. The bar chart shows the mean % positive cells and error bars representing SEM across three independent replicates. (D) WT H9 cells were transfected with dCas9 activator modRNA and NKX2.5 dgRNA on day 0. Cells were collected for RNA extraction and qPCR analysis of NKX2.5, 48 hours after transfection. Relative expression of NKX2.5 is shown in transfected cells vs untransfected cells. (E-G) WT H9 cells were transfected with dCas9 activator modRNA and PDX1 dgRNA on day 0. On day 5, cells were immunostained for canonical pluripotency markers OCT4, NANOG, SOX2, and TRA-1-60 and analyzed by either immunofluorescent imaging (E) or flow cytometry (F-G). Scale bar, 100 μm. Representative flow cytometry plots showing day 5 expression of OCT4 and TRA-1-60. (G) Quantification of (F) with error bars representing SEM across three independent replicates. (H) Diagram of our two-step approach for deriving reporter hPSCs using either plasmid or modRNA transfection to deliver CRISPR components. (I-J) NKX2.5-nEGFP reporter cells were generated from WT H9 cells as shown in (H). Cells were analyzed for GFP expression by flow cytometry, 48 hours after activating NKX2.5 by either plasmid-based or modRNA-based approaches. (I) Representative flow cytometry plots showing GFP expression across each of the four combinations. (J) Quantification of (I) with error bars representing SEM across 3 independent replicates. p < 0.05 (*), p < 0.01 (**), p < 0.001 (***).

While activation with our modSAM initially induced morphological changes in cultured hPSCs, we observed that cells reverted to the typical hPSC morphology 4 to 5 days later. Consequently, we decided to evaluate whether hPSCs maintain pluripotent state 5 days post-transfection by analyzing the expression of pluripotency markers. Nearly 100% of day 5 cells were positive for OCT4, NANOG and SOX2 based on immunostaining assays (**Fig 1E, S1A**). This was confirmed by flow cytometry analysis of day 5 cells which showed high levels of OCT4 and TRA-1-60 expression similar to wild-type (WT) hPSCs (**Fig 1F-G, S1B-E**). Considering the transient nature of the activation and the absence of irreversible differentiation in the modSAM transfected cells, we hypothesized that our modSAM system might serve as a selection tool for purifying correctly targeted cells after Cas9-mediated knockin of a fluorescent protein to the lineage-specific genes. At present, the selection for successful knock-in of lineage-specific genes necessitates a laborious process involving the selection and validation of single-cell clones through genomic PCR and sequencing.

In order to test our proposed strategy and show the efficacy of our modRNA-based approach against traditional plasmid-based delivery methods, we designed an experiment to create NKX2.5-nEGFP (EGFP with nuclear localization signal sequence) reporter cells using parallel methods of plasmid-based or modRNA-based delivery of Cas9 and dCas9 activator (**Fig 1H**).

Our MAGIK strategy employs a two-step approach: first, we use a Cas9-2A-p53DD modRNA (iCas9 modRNA), a sgRNA targeting the stop codon region (KI sgRNA), and a mini-donor plasmid (no drug selection cassette) to insert the fluorescent protein. The iCas9 modRNA was used because it can generate higher gene editing efficiency as compared to Cas9 modRNA (**Fig. S1F**). Subsequently, we transiently activate the target gene using modSAM to enable sorting of the positive cells on day 2 post-transfection.

For our initial screen, we performed each step using either plasmid-based or modRNA-based delivery, yielding a total of 4 possible combinations. We analyzed the efficiency of each combination via flow cytometry of day 2 activated cells. Consistent with our initial hypothesis, our completely modRNA-based approach significantly outperformed all other combinations, showing a nearly **45-fold** improvement over an entirely plasmid-based approach (**5.77% ± 0.27 vs 0.13% ± 0.03**) (**Fig 1I-J**). We validated our results in a second gene via generation of PDX1-EGFP H9 reporter cells. We observed a **15-fold** improvement with our entirely modRNA-based strategy over the traditional plasmid-based delivery of CRISPR components (**Fig S1G-H**).

### Generation of NKX6.1-nEGFP reporter line via MAGIK

Considering the effectiveness of our approach in achieving significantly higher proportions of successfully edited cells, we chose to utilize our MAGIK method to create an NKX6.1-nEGFP H9 reporter line (**Fig 2A**). NKX6.1 is an important marker for pancreatic progenitors and beta cells^18,19^. We cloned an NKX6.1-nEGFP mini-donor plasmid (no drug-selection cassette), based on the traditional NKX6.1 donor plasmid previously reported^20^. Additionally, we synthesized a KI sgRNA targeting the stop codon region of NKX6.1 based on a published sequence^20^ (**Fig 2B**).

**Figure 2.**
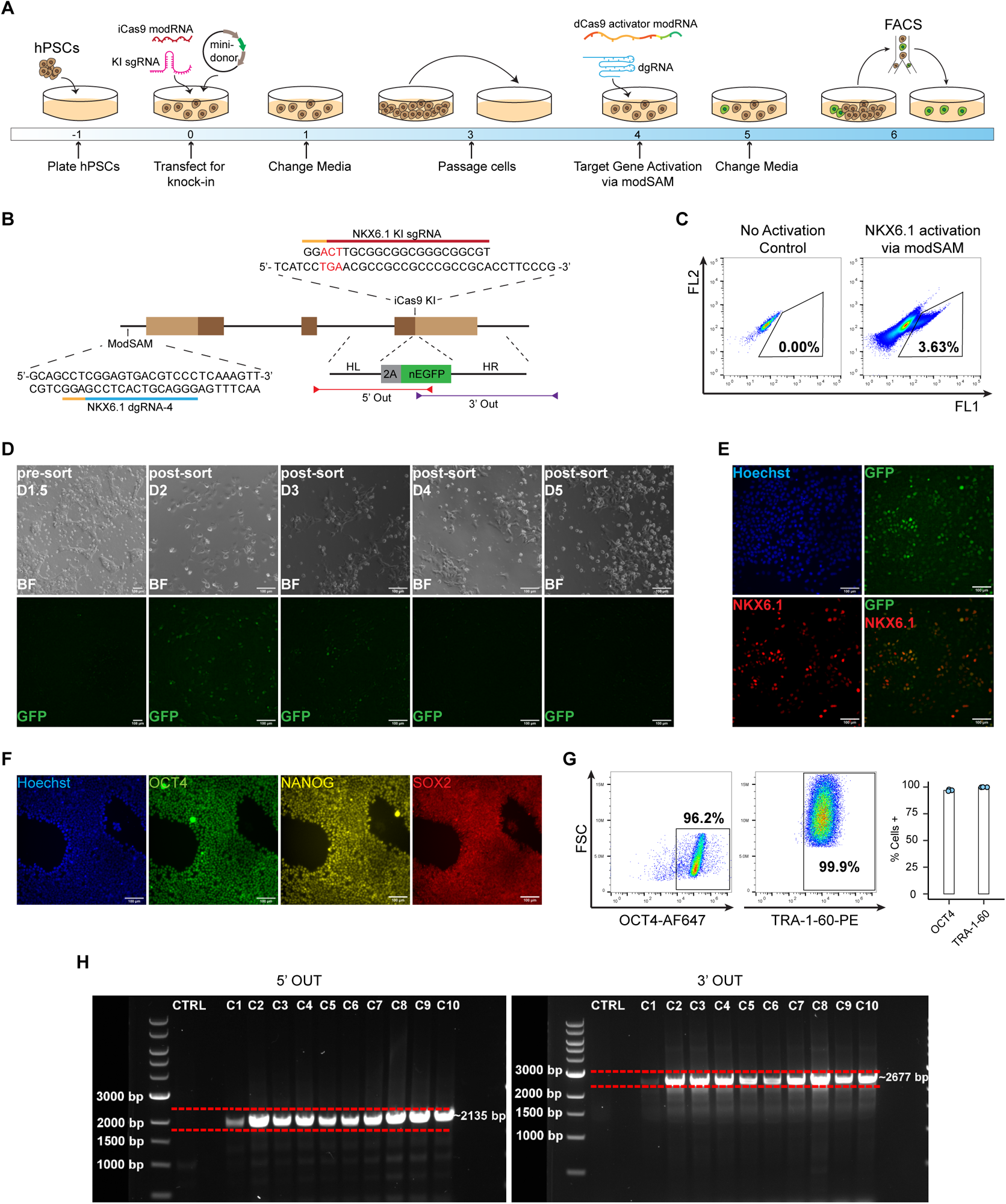
MAGIK for generating H9 NKX6.1-nEGFP reporter cells. (A) Schematic diagram showing timeline for MAGIK approach for generating fluorescent reporter hPSCs. (B) Diagram showing knock-in and activation strategy for generating NK6.1-nEGFP reporter cell line. (C) WT H9 cells were transfected with iCas9 modRNA, NKX6.1 KI sgRNA, and NKX6.1-nEGFP mini-donor plasmid. Cells were expanded in TeSR for several days before being passaged into multiple wells of a 12-well plate. The mixed knock-in cells were transfected with dCas9 activator modRNA and NKX6.1 dgRNA. EGFP^+^ cells were sorted via FACS 48 hours after transfection and replated into a single well of a 48-well plate. Representative plot showing percentage of EGFP positive cells sorted from total population. (D) Representative brightfield and fluorescent images of knock-in reporter cells before and after sorting until day 5 post-transfection. Scale bar, 100 μM. (E) Post-sort H9 NKX6.1-nEGFP reporter cells were transfected with dCas9 activator modRNA and NKX6.1 dgRNA. On day 2 following transfection, cells were fixed and stained for NKX6.1 (Red). Representative fluorescent images showing co-localization of anti-NKX6.1 signal and GFP. Scale bar, 100 μM. (F-G) Post-sort H9 NKX6.1-nEGFP reporter cells were stained for pluripotency markers OCT4, NANOG, SOX2 and TRA-1-60 and analyzed by either immunofluorescent imaging (F) or flow cytometry (G). Scale bar, 100 μm. Representative flow cytometry plots and quantification of OCT4 and TRA-1-60 expression in post-sort H9 NKX6.1-nEGFP reporter cells. Error bars represent SEM across three replicates. (H) PCR genotyping of ten single-cell derived clones from post-sort H9 NKX6.1-nEGFP reporter cells. The expected band from each set of probes is highlighted by a pair of red dashed lines (5’ OUT: 2135 bp; 3’ OUT: 2677 bp).

H9 cells were initially transfected with iCas9 modRNA, NKX6.1 KI sgRNA, and NKX6.1-nEGFP mini-donor plasmid. After a cultivation period, cells were passaged and transfected with our modSAM system (dCas9 activator modRNA + NKX6.1-dgRNA4). Two days post-transfection, EGFP^+^ cells were sorted and replaced. The purity of EGFP^+^ cells was 3.63%, compared to 0% in the untransfected population (**Fig. 2C**). We confirmed EGFP expression via fluorescent microscopy on the day of sorting (**Fig. 2D**). Subsequently, all cells were EGFP^-^ on day 5 post-transfection (**Fig. 2D**).

To assess whether the sorted cells were indeed NKX6.1-nEGFP knockin cells, we transfected the sorted cells via modSAM and immunostained the cells using a NKX6.1 antibody. Our data showed co-localization of EGFP with anti-NKX6.1 antibody signal (**Fig 2E**). To study whether the sorted cells retained an undifferentiated state, we assessed the cells via immunostaining of pluripotency makers. Our data showed the sorted cells exhibited nearly 100% expression of OCT4, NANOG, and SOX2 (**Fig 2F**), with flow cytometry revealing similar expression levels of OCT4 and TRA-1-60 to WT H9 cells (**Fig 2G, S1B**).

From the sorted cell population, we derived single cell clones and extracted genomic DNA (gDNA). PCR on the gDNA with our 5’ Out and 3’ Out primers (**Fig. 2B**) confirmed successful integration of EGFP at the expected site in all clonal populations (**Fig. 2H**).

### Generation of SOX17-nEGFP reporter line via MAGIK

SOX17 gene encodes a member of the SOX (SRY-related HMG-box) family of transcription factors. It plays a crucial role in embryo development and serves as a marker for definitive endoderm cells^21,22^. Additionally, SOX17 specifies human primordial germ cell^23^ and blood cell fate^24^. To generate a SOX17-nEGFP H9 reporter line using our MAGIK method, we cloned a SOX17-nEGFP mini-donor plasmid with the appropriate left and right homology arms (**Fig. 3A**), and synthesized a SOX17 KI sgRNA based on a published sequence^25^. The knock-in and sorting of EGFP^+^ cells were performed after modSAM transfection (**Fig. 2A**). During sorting, the purity of EGFP^+^ cells was 0.51% for transfected cells, compared to 0% in the untransfected population (**Fig. 3B**). After sorting, EGFP^+^ cells were seeded into a well of the 48-well plate, and EGFP expression was monitored daily until no longer detectable (**Fig. 3C**). Similar to the sorted NKX6.1-nEGFP reporter, all sorted SOX17-nEGFP cells exhibited EGFP expression under fluorescent microscopy (**Fig. 3C**). In addition, the sorted cells lost EGFP expression starting from day 4 post-sorting (**Fig. 3C**).

**Figure 3.**
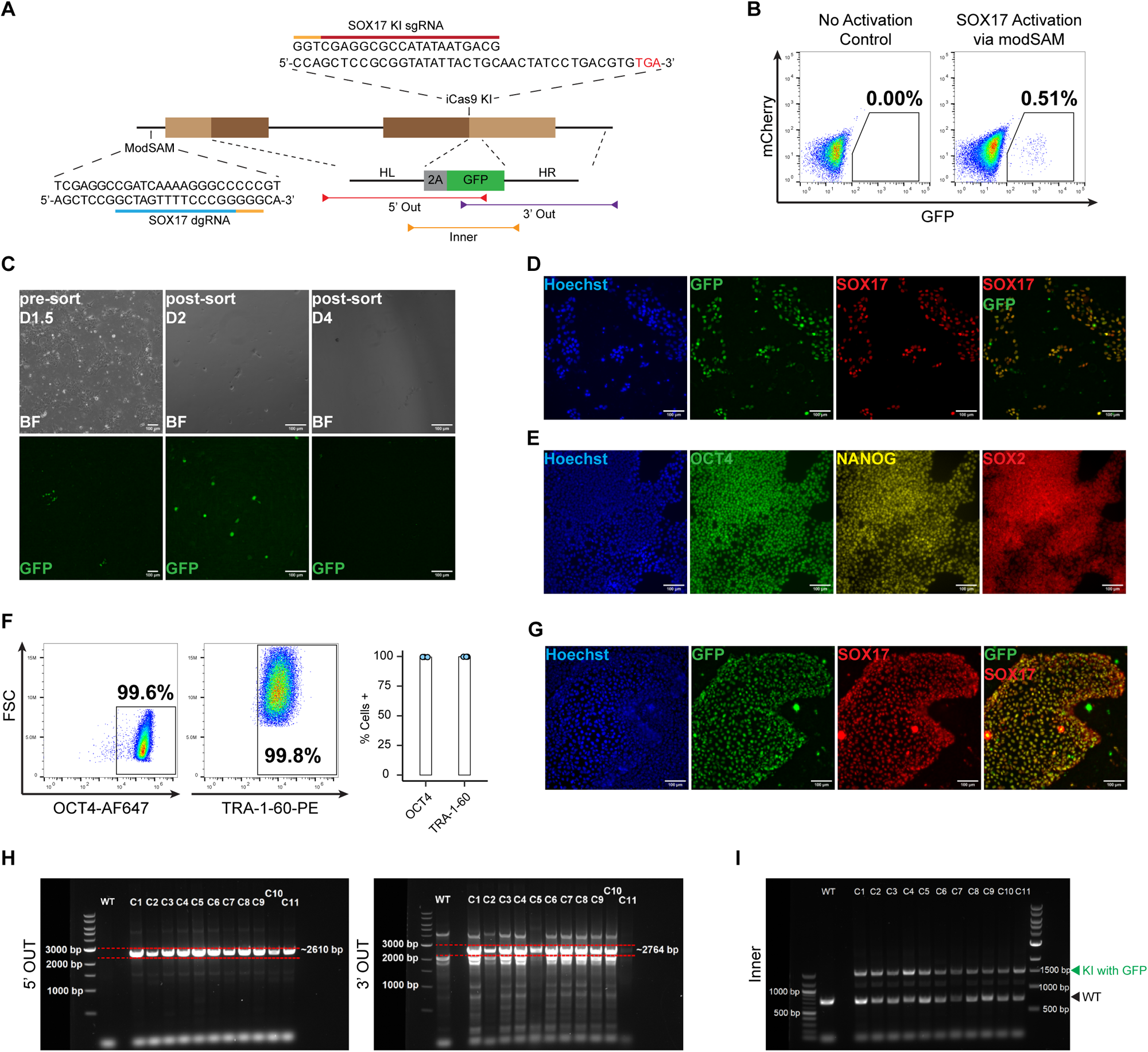
MAGIK for generating H9 SOX17-nEGFP reporter cells. (A) Diagram showing knock-in and activation strategy for generating SOX17-nEGFP reporter cell line. (B) WT H9 cells were transfected with iCas9 modRNA, SOX17 KI sgRNA, and SOX17-nEGFP mini-donor plasmid. Cells were expanded in TeSR for several days before being passaged into multiple wells of a 12-well plate. The mixed knock-in cells were transfected with dCas9 activator modRNA and SOX17 dgRNA. EGFP^+^ cells were sorted via FACS 48 hours after transfection and replated into a single well of a 48-well plate. Representative plot showing percentage of EGFP positive cells sorted from total population. (C) Representative brightfield and fluorescent images of knock-in reporter cells before and after sorting until day 4 post-transfection. Scale bar, 100 μM. (D) Post-sort H9 SOX17-nEGFP reporter cells were transfected with dCas9 activator modRNA and SOX17 dgRNA. On day 2 following transfection, cells were fixed and stained for SOX17 (Red). Representative fluorescent images showing co-localization of anti-SOX17 signal and EGFP. Scale bar, 100 μM. (E-F) Post-sort H9 SOX17-nEGFP reporter cells were stained for pluripotency markers OCT4, NANOG, SOX2 and TRA-1-60 and analyzed by either immunofluorescent imaging (E) or flow cytometry (F). Scale bar, 100 μm. Representative flow cytometry plots and quantification of OCT4 and TRA-1-60 expression in post-sort H9 SOX17-nEGFP reporter cells. Error bars represent SEM across three replicates. (G) H9 SOX17-nEGFP reporter cells were differentiated to definitive endoderm. Day 4 differentiated cells were fixed and stained for SOX17 (Red). Representative fluorescent images showing co-localization of anti-SOX17 signal and GFP. Scale bar, 100 μm. (H) PCR genotyping of eleven single-cell derived clones from post-sort H9 SOX17-nEGFP reporter cells. The expected band from each set of probes is highlighted by a pair of red dashed lines (5’ OUT: 2610 bp; 3’ OUT: 2764 bp). (I) PCR genotyping of single-cell derived clones using Inner primers to distinguish between monoallelic and biallelic knockin (WT: 814 bp; KI with GFP: 1615 bp)

To validate whether the sorted cells were indeed SOX17 knockin cells, we transfected the sorted cells via modSAM using a validated SOX17 dgRNA. Our data showed co-localization of EGFP with anti-SOX17 antibody signal (**Fig. 3D**). To assess whether the sorted cells retained an undifferentiated state, we characterized the cells via immunostaining of pluripotency makers. Our data showed the sorted cells exhibited nearly 100% expression of OCT4, NANOG, and SOX2 (**Fig. 3E**), with flow cytometry revealing nearly 100% expression of OCT4 and TRA-1-60 (**Fig. 3F**). We then differentiated the sorted cells into definitive endoderm (DE) cells using our small-molecule DE protocol^22^. On day 4 of differentiation, the differentiated cells were fixed and immunostained with a SOX17 antibody. Our data revealed that the EGFP expression co-localized with the anti-SOX17 immunofluorescence signal (**Fig. 3G**). In order to provide further validation of our sorted SOX17-nEGFP H9 reporter line, we performed PCR genotyping of gDNA extracted from 11 single-cell derived clones using our 5’ Out and 3’ Out primers. Running the PCR product on DNA gels revealed successful integration of the reporter construct in all clonal populations (**Fig. 3H**). Additionally, PCR amplification of gDNA using our Inner primers revealed that all single-cell derived clones were heterozygous for the fluorescent reporter (**Fig. 3I**).

### MAGIK for generation of NKX2.5-nEGFP reporter

NKX2.5 is transcription factor that plays a critical role in the specification of cardiac fate, and defects in this gene are associated with congenital heart diseases^26^. For development of an NKX2.5-nEGFP H9 reporter, we chemically synthesized a mini-donor plasmid with left and right homology arms (**Fig. 4A**). Additionally, we designed and synthesized a NKX2.5 KI sgRNA. H9 cells were transfected with iCas9 modRNA, NKX2.5 KI sgRNA and NKX2.5-nEGFP mini-donor plasmid, followed by modSAM transfection. Two days after modSAM transfection, EGFP+ cells (**5.9%**) were sorted and seeded onto a pre-coated well of the 48-well plate (**Fig. 4B**). EGFP expression was monitored via fluorescent microscopy until EGFP expression had completely ablated on the fourth day post modSAM transfection (**Fig. 4C**). Similar to NKX6.1 and SOX17 knockin cells, we validated our sorted NKX2.5-nEGFP cells via modSAM transfection and immunostaining for NKX2.5 expression. Fluorescent microscopy of immunostained cells showed co-localization of EGFP signal with anti-NKX2.5 antibody signal (**Fig 4D**). Additionally, sorted reporter cells showed nearly 100% expression of pluripotency markers OCT4, NANOG, and SOX2 as determined by immunostaining (**Fig. 4E**). Flow cytometry of reporter cells stained for OCT4 and TRA-1-60 showed expression equivalent to WT H9 cells (**Fig. 4F, S1B**). NKX2.5-nEGFP reporter cells were differentiated into cardiomyocytes using our previously reported GiWi protocol^27,28^. Cells assayed on day 10 of differentiation were 74.0% ± 0.6% EGFP^+^ according to flow cytometry (**Fig. 4G**). Immunostaining of day 10 cells with anti-NKX2.5 antibody revealed that EGFP signal co-localized with the anti-NKX2.5 antibody (**Fig. 4H**). We derived 12 single-cell clonal lines from the sorted H9 NKX2.5-nEGFP reporter and extracted gDNA for PCR genotyping. Based on the results from our 5’ Out and 3’ Out primers we validated successful integration of the reporter construct across all single-cell clonal populations tested (**Fig. 4I-J**). Additionally, using our Inner primers we determined that 2 of the 12 clonal populations had bi-allelic integration of the fluorescent reporter (**Fig. 4K**).

**Figure 4.**
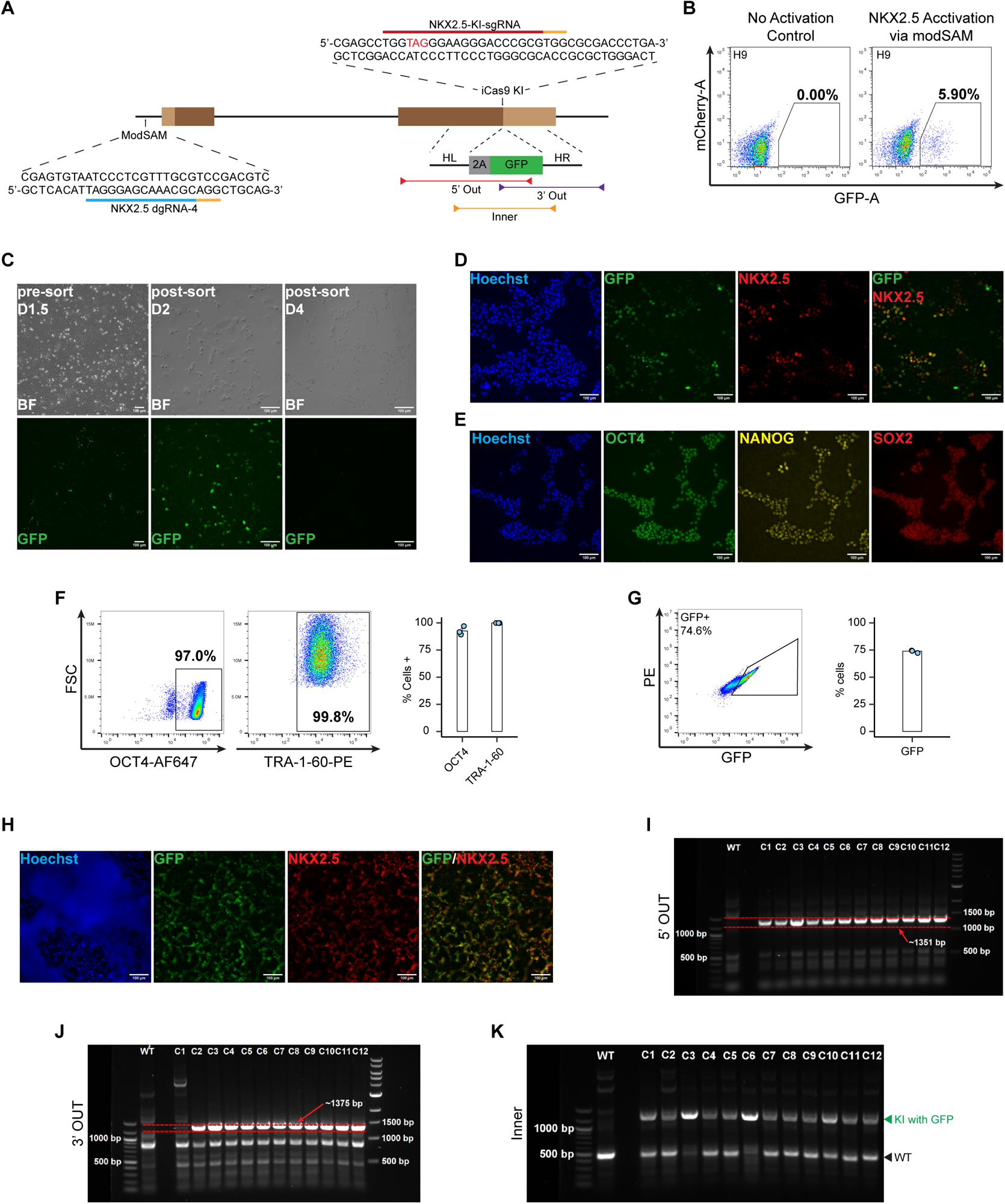
MAGIK for generating H9 NKX2.5-nEGFP reporter cells. (A) Diagram showing knock-in and activation strategy for generating NKX2.5-nEGFP reporter cell line. (B) WT H9 cells were transfected with iCas9 modRNA, NKX2.5 KI sgRNA, and NKX2.5-nEGFP mini-donor plasmid. Cells were expanded in TeSR for several days before being passaged into multiple wells of a 12-well plate. The mixed knock-in cells were transfected with dCas9 activator modRNA and NKX2.5 dgRNA. EGFP^+^ cells were sorted via FACS 48 hours after transfection and replated into a single well of a 48-well plate. Representative plot showing percentage of GFP positive cells sorted from total population. (C) Representative brightfield and fluorescent images of knock-in reporter cells before and after sorting until day 4 post-transfection. Scale bar, 100 μm. (D) Post-sort H9 NKX2.5-nEGFP reporter cells were transfected with dCas9 activator modRNA and NKX2.5 dgRNA. On day 2 following transfection, cells were fixed and stained for NKX2.5 (Red). Representative fluorescent images showing co-localization of anti-NKX2.5 signal and GFP. Scale bar, 100 μm. (E-F) Post-sort H9-NKX2.5nEGFP reporter cells were stained for pluripotency markers OCT4, NANOG, SOX2 and TRA-1-60 and analyzed by either immunofluorescent imaging (E) or flow cytometry (F). Scale bar, 100 μm. Representative flow cytometry plots and quantification of OCT4 and TRA-1-60 expression in post-sort H9 NKX2.5-nEGFP reporter cells. Error bars represent SEM across three replicates. (G-H) H9 NKX2.5-nEGFP reporter cells were seeded on Matrigel-coated plates and differentiated into cardiomyocytes using our GiWi protocol. Day 10 cells were either (G) collected for live-cell flow cytometry or (H) fixed and stained for NKX2.5 for immunofluorescent imaging. (G) Representative flow cytometry plot and quantification of EGFP expression. Error bars represent SEM across three replicates. (H) Scale bar, 100 μm. (I-J) PCR genotyping of twelve single-cell derived clones from post-sort H9 NKX2.5-nEGFP reporter cells. The expected band from each set of probes is highlighted by a pair of red dashed lines (5’ OUT: 1351 bp; 3’ OUT: 1375 bp). (K) PCR genotyping of single-cell derived clones using Inner primers to distinguish between mono-allelic and bi-allelic integration of the fluorescent reporter (WT: 499 bp; KI with GFP: 1303 bp)

### MAGIK for generation of PDX1-EGFP reporter line in multiple hPSC lines

Having demonstrated the effectiveness of MAGIK in creating fluorescent reporter lines spanning various genes within H9 cells, we sought to authenticate our methodology across a spectrum of hPSC lines. To accomplish this objective, we elected to produce PDX1-EGFP reporters utilizing multiple hPSC lines, including H9, H1, IMR90 iPSCs, and 6-9-9 iPSCs.

PDX1 functions as a pivotal regulator in pancreas development^29^. Throughout the differentiation of hPSCs into pancreatic lineages, PDX1 becomes expressed within pancreatic progenitors and beta cells^22,30^. In the context of PDX1 knockin, we leveraged our iCas9 modRNA in conjunction with a PDX1-EGFP mini-donor plasmid^12^ and a PDX1 KI sgRNA (**Fig 5A**). To effectively isolate cells that had undergone successful targeting, we transfected KI cells with our modSAM system, utilizing our validated PDX1 dgRNA. Encouragingly, the MAGIK method yielded positive outcomes across all four cell lines. Two days after modSAM transfection, the proportions of EGFP^+^ cells were 0.44% (H9 hESC), 1.41% (IMR90 iPSC), 0.76% (H1 hESC), and 1.76% (6-9-9 iPSC) (**Fig. 5B, S2A, S3A, and S4A**).

**Figure 5.**
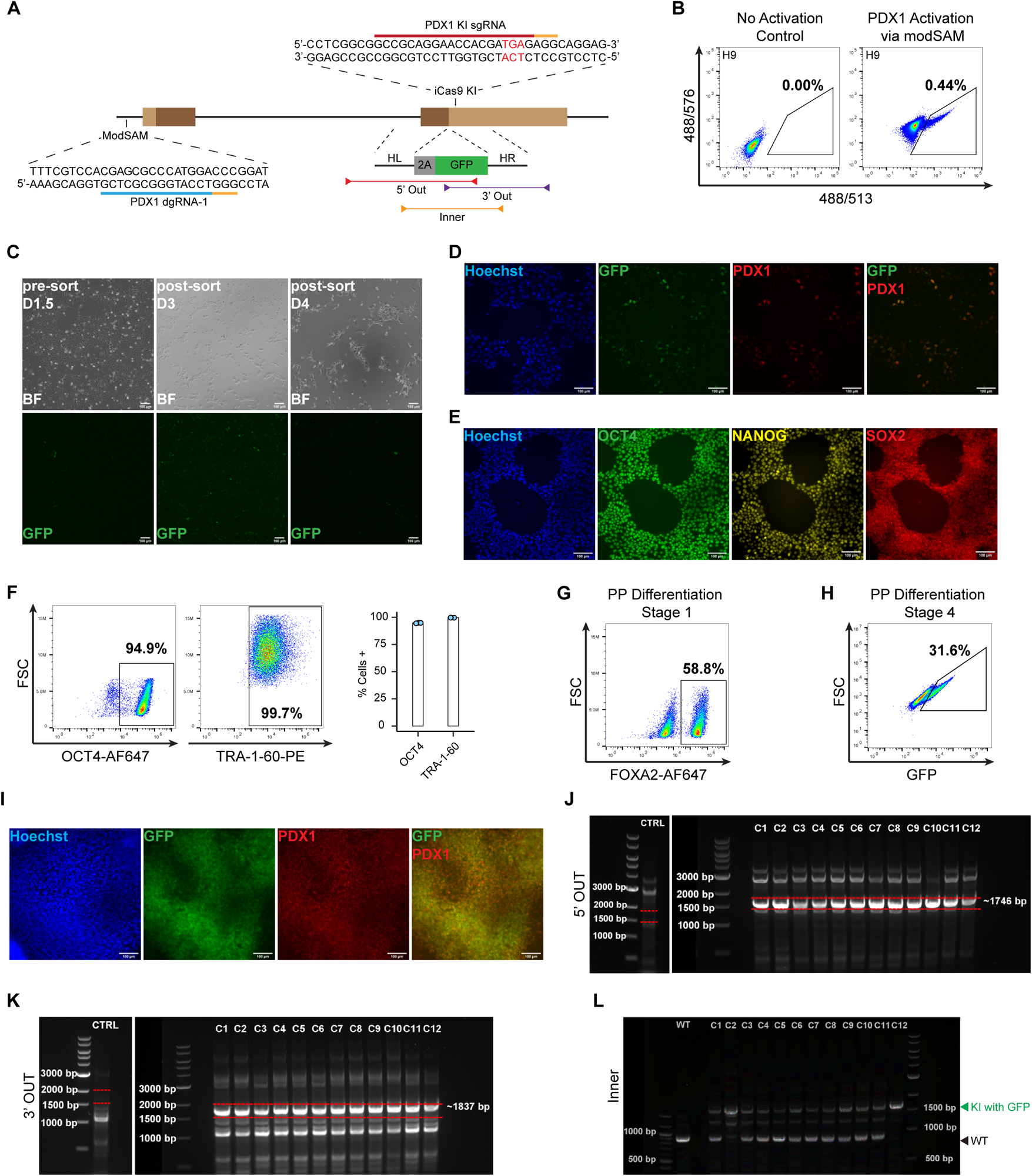
MAGIK for generating PDX1-EGFP reporter in multiple hPSC lines. (A) Diagram showing knock-in and activation strategy for generating PDX1-EGFP reporter cell line. (B) WT H9 cells were transfected with iCas9 modRNA, PDX1 KI sgRNA, and PDX1-EGFP mini-donor plasmid. Cells were expanded in TeSR for several days before being passaged into multiple wells of a 12-well plate. The mixed knock-in cells were transfected with dCas9 activator modRNA and PDX1 dgRNA. EGFP+ cells were sorted via FACS 48 hours after transfection and replated into a single well of a 48-well plate. Representative plot showing percentage of EGFP positive cells sorted from total population. (C) Representative brightfield and fluorescent images of knock-in reporter cells before and after sorting until day 4 post-transfection. Scale bar, 100 μm. (E) Post-sort H9 PDX1-EGFP reporter cells were transfected with dCas9 activator modRNA and PDX1 dgRNA. On day 2 following transfection, cells were fixed and stained for PDX1 (Red). Representative fluorescent images showing co-localization of anti-PDX1 signal and EGFP. Scale bar, 100 μm. (E-F) Post-sort H9 PDX1-EGFP reporter cells were stained for pluripotency markers OCT4, NANOG, SOX2 and TRA-1-60 and analyzed by either immunofluorescent imaging (E) or flow cytometry (F). Scale bar, 100 μm. Representative flow cytometry plots and quantification of OCT4 and TRA-1-60 expression in post-sort H9 PDX1-EGFP reporter cells. Error bars represent SEM across three replicates. (G-I) H9 PDX1-EGFP reporter cells were differentiated to pancreatic progenitors (PP) using the GiBi protocol. (G) Day 4 cells were collected and analyzed for FOXA2 expression by flow cytometry. (H) Day 14 cells were collected and analyzed for EGFP expression by flow cytometry. (I) Day 14 cells were fixed and stained for PDX1 (Red). Representative fluorescent images showing colocalization of anti-PDX1 signal and GFP. Scale bar, 100 μm. (J-K) PCR genotyping of twelve single-cell derived clones from post-sort H9 PDX1-EGFP reporter cells. Expected band from each set of probes is highlighted by a pair of red dashed lines (5’ OUT: 1746 bp; 3’ OUT: 1837 bp). (L) PCR genotyping of single-cell derived clones using Inner primers to distinguish between monoallelic and biallelic knock in (WT: 885 bp; KI with GFP: 1659 bp)

Subsequently, EGFP^+^ sorted cells were seeded into individual wells of a 48-well plate. Upon observation with a fluorescent microscope, it was confirmed that the attached cells were indeed expressing EGFP (**Fig. 5C, S2B-C, S3B-C, and S4B-C**). As expected, all sorted cells ceased EGFP expression within one week of plating (**Fig. 5C, S2B-C, S3B-C, and S4B-C**). Following expansion, the sorted cells underwent transfection with our modSAM system to activate PDX1 and EGFP expression. Immunostaining analyses showed co-localization between the EGFP signal and anti-PDX1 immunofluorescence signal (**Fig. 5D, S2D, S3D, and S4D**). Moreover, the sorted cells consistently displayed nearly 100% expression of the pluripotency markers OCT4, NANOG, and SOX2, as determined through immunostaining across all tested cell lines (**Fig. 5E, S2E, S3E, and S4E**). Additionally, the percentage of cells positive for OCT4 and TRA-1-60, as determined by flow cytometry, exhibited similarities between the sorted reporter lines and WT H9 cells (**Fig. 5F, S2F, S3F, and S4F**). These findings collectively signify the feasibility of employing the MAGIK method to generate lineage-specific reporters within diverse hPSC populations.

Next, we differentiated PDX1-EGFP H9 cells into pancreatic β-cell using our small-molecule-based protocol^22^. At the end of stage 1, we obtained 58.8% FOXA2^+^ DE cells (**Fig. 5G**). Additionally, at the end of stage 4, 31.6% of PDX1-EGFP reporter cells were EGFP^+^, indicating that our knockin reporter cells can accurately report the expression of PDX1 (**Fig 5H**). We further performed immunostaining of stage 4 cells using a PDX1 antibody. Our data showed co-localization of EGFP expression with the anti-PDX1 signal (**Fig 5I**).

We derived multiple single cell clonal populations from our sorted H9, H1 and IMR90 PDX1-EGFP reporters and extracted gDNA for PCR analysis. Using our 5’ Out and 3’ Out probes we showed successful amplification of the expected bands, demonstrating successful integration of the fluorescent reporter at the target site across all clonal populations in all three cell lines (**Fig 5J, 5K, S2G, S3G**). Additionally, PCR amplification of the gDNA using the Inner primers indicated that 5 clonal lines out of the 32 clonal populations tested across the three cell lines (15.6%) were homozygous for reporter integration (**Fig. 5L, S2H, S3H).**

Overall, we showed our MAGIK method can be used to derive fluorescent reporter lines across multiple lineage-specific genes and demonstrated the robustness of MAGIK across multiple stem cell lines.

## DISCUSSION

The ability to perform knockin in hPSCs is vital for the advancement of stem cell research and therapy. The introduction of fluorescent proteins into lineage-specific marker genes within hPSCs has revolutionized the isolation of desired differentiated cells. However, conventional knock-in methodologies have long grappled with challenges like low efficiency and intricate protocols. The necessity of integrating a drug-selection cassette into the donor plasmid, coupled with the subsequent requirement to excise the drug selection component post-successful knockin, has elongated the knock-in process significantly.

Scientists have developed various strategies to enhance the efficiency of precise sequence replacement or insertion via homology-directed repair (HDR). By strategically designing single-stranded DNA (ssDNA) donors with optimal length complementary to the strand initially released by Cas9, researchers can boost HDR rates in human cells, achieving rates as high as 60% (^31^). However, synthesizing long ssDNA poses relative difficulty. For employing double-stranded DNA (dsDNA) donors, researchers have discovered that modifying dsDNA, such as using specially designed 3’-overhang dsDNA donors containing 50-nt homology arms^32^ or employing sgRNA-PAM sequence-flanked dsDNA as donors^33^, can significantly enhance HDR efficiency. These specialized dsDNA donors have not been demonstrated in hPSCs and these special donors present greater production challenges compared to traditional plasmid donors. Additionally, researchers have explored small-molecule inhibitors to enhance HDR efficiency. They identified AZD7648, an inhibitor of DNA-dependent protein kinase catalytic subunit (DNA-PKcs), a crucial protein in the alternative repair pathway of non-homologous end joining (NHEJ), which can substantially increase HDT efficiency (up to 50-fold)^34^. Simultaneously inhibiting DNA-PK and PolL can further improve HDR efficiency and the precision of genome editing^35^. Nevertheless, despite these multiple approaches to enhance HDR efficiency, it’s important to note that HDR efficiency is not guaranteed to reach 100%. Consequently, achieving precise integration of a fluorescent protein into a silenced lineage-specific marker gene still necessitates single cell cloning, a step that considerably delays the knockin process.

To address these challenges, we introduce the MAGIK method. This novel strategy enhances knockin efficiency by employing an iCas9 modRNA and facilitates target cell isolation with a modSAM system. Our method simplifies the process of isolating successful knockin cells, providing a markedly efficient and expedited alternative to conventional methods. The MAGIK method stands out for its efficiency and simplicity. It starts with the use of an iCas9 modRNA to significantly enhance knockin efficiency. The subsequent step employs a dCas9 activator modRNA to temporarily activate the expression of lineage-specific genes and their associated fluorescent proteins. This maneuver simplifies cell isolation through FACS, negating the need for selection and validation of single cell-derived clones. The MAGIK results showed a 45-fold improvement over an entirely plasmid-based approach. We demonstrate the robustness of MAGIK across multiple lineage-specific genes (*NKX6.1, SOX17, NKX2.5,* and *PDX1*) and four hPSC lines. The method’s ability to derive relatively pure fluorescent reporter cells without the use of drug selection cassettes and arduous genomic PCR screening makes it a significant advancement in the field.

In summary, the MAGIK method represents a significant step forward in the field of stem cell research. Its efficiency, simplicity, and applicability across various genes and stem cell lines make it a promising tool for developing lineage-specific reporters. The future applications of MAGIK could extend to various areas of biomedical research, offering new avenues for understanding and treating human diseases.

## RESOURCE AVAILABILITY

### Lead contact

Further information and requests for resources and reagents should be directed to and will be fulfilled by the lead contact, Dr. Xiaojun Lance Lian.

### Materials availability

All plasmids generated from this paper will be available at addgene.

### Data and code availability

The published article includes all the dataset generated during this study. This paper does not report original code. Any additional information required to re-analyze the data reported in this paper is available from the lead contact upon request.

## EXPERIMENTAL MODEL AND SUBJECT DETAILS

### Cell lines

Four pluripotent cell lines, H9, H1, IMR90C4 and 6-9-9 (**Table S1**) were used for this study. These lines were obtained from WiCell Research Institute. All cell culture experiments involving human pluripotent stem cell lines were approved by the Embryonic Stem Cell Oversight Committee at the Pennsylvania State University and carried out in accordance with the approved guidelines.

## METHOD DETAILS

### Maintenance of hPSCs

hPSCs were maintained on iMatrix-511 (Iwai North America) coated plates in mTeSR plus (TeSR) medium (STEMCELL Technologies). Cells were regularly passaged when they reached 80-90% confluency, usually 3-4 days after the previous passage. For passaging, cell medium was aspirated and 1 ml of Accutase (Innovative Cell Technologies) was added to each well. Cells were incubated at 37°C, 5% CO_2_ for 5 to 10 minutes. Dissociated cells were transferred to excess DMEM at a 1:2 (vol/vol) ratio and centrifuged at 1000 rpm for 4 minutes. New wells were precoated with 0.75 μg/ml iMatrix-511 and incubated at 37°C, 5% CO_2_ for 10 minutes. After centrifugation, cell pellet was resuspended in TeSR with 5 μM Y-27632 (Selleck Chemicals). 10,000-20,000 cells/cm^2^ were seeded onto iMatrix-511 coated wells. For regular maintenance, hPSCs were cultured in the six-well plates.

### ModRNA synthesis

To clone desired DNA fragments into modRNAc1 plasmid, VP64dCas9VP64-T2A-MPH and Cas9-2A-p53DD was PCR amplified from appropriate plasmids (**Table S2, S3**). The PCR product was run on a 1% Agarose gel and the band at the appropriate size was excised and the DNA extracted using the Zymoclean Gel DNA Recovery kit (Zymo Research). Purified insert DNA was cloned into the linearized modRNAc1 plasmid using the In-Fusion Cloning Kit (Takara Bio). The DNA template for modRNA synthesis was PCR amplified from the successfully cloned modRNAc1 plasmid followed by PCR purification using DNA Clean & Concentrator-5 (Zymo Research). ModRNA was synthesized from the PCR DNA template via *in vitro* transcription (IVT) using the MEGAscript T7 Transcription kit (ThermoFisher) supplemented with 12.5 mM ATP, 12.5 mM GTP, 12.5 mM CTP, 12.5 mM N1-methyl-pseudo-UTP (TriLink Biotechnologies), and 10 mM CleanCap AG (TriLink Biotechnologies). The IVT reaction product was treated with DNase I to remove DNA template and then purified using the Monarch RNA clean-up kit (NEB). RNA concentration was measured using a NanoDrop (ThermoFisher).

### sgRNA synthesis

sgRNA was synthesized using the EnGen sgRNA synthesis kit (NEB). Target specific oligos (**Table S4**) were ordered from Integrated DNA Technologies using the following template: *TTCTAATACGACTCACTATA***G**(N_20_)**GTTTTAGAGCTAGA**. The IVT reaction was assembled based on the EnGen sgRNA synthesis kit’s recommendations. The sgRNA was purified using an RNA Clean & Concentrator-5 kit (Zymo Research). RNA concentration was measured using a NanoDrop.

### dgRNA synthesis

Spacer sequences of dgRNA for gene activation of NKX6.1 and PDX1 were selected using the integrated CRISPR tool in Benchling (**Table S5, S6**). The spacer sequence for NKX2.5 activation was selected using the ChopChop online tool^36^. The spacer sequence for SOX17 activation was previously described by Kearns et al.^17^. The DNA template for dgRNA synthesis was assembled via PCR of variable dgRNA-Oligo1, dgRNA-Oligo2 and dgRNA-Oligo3. The PCR product was run on a 2% Agarose gel and the band at 150 bp was excised and then purified using the Zymoclean Gel DNA Recovery kit (Zymo Research). The dgRNA was synthesized via IVT using the MEGAscript T7 Transcription kit supplemented with 10mM ATP, 10mM GTP, 10mM CTP, and 10mM UTP. The IVT reaction product was treated with DNase I to remove the DNA template and then purified using the Monarch RNA clean-up kit (NEB). dgRNA concentration was measured using a NanoDrop.

### Generation of mini-donor plasmids

To generate a mini-donor plasmid (no drug selection cassette) for NKX6.1, the left homology arm-2A-nEGFP, and the right homology arm were PCR amplified from the NKX6.1-2A-nEGFP-PGK-Puro donor plasmid^20^, which was provided by Dr. Haisong Liu and Dr. Kai Wang. These two PCR fragments were cloned into the empty backbone derived from the PDX1-2A-EGFP mini-donor plasmid. PDX1-2A-eGFP mini-donor plasmid was obtained from Addgene (#66964). For SOX17 targeting, we cloned a mini-donor plasmid (SOX17-2A-nEGFP) containing: i) a left homology arm containing a 1.9 kb sequence immediately upstream of the SOX17 stop codon; ii) a 2A-nEGFP cassette; and iii) a right homology arm containing a 1.9 kb sequence downstream of the SOX17 stop codon. The left and right homology arms were PCR amplified from genomic DNA of hPSCs using appropriate primers. NKX2.5-2A-nEGFP mini-donor plasmid was chemically synthesized containing: i) a left homology arm containing a 0.5 kb sequence immediately upstream of the NKX2.5 stop codon; ii) a 2A-nEGFP cassette; and iii) a right homology arm containing a 0.5 kb sequence downstream of the NKX2.5 stop codon. All cloning was performed using the In-Fusion Cloning kit (Takara Bio) and ligated plasmids were transformed into chemically competent *Stbl3* E. coli cells.

### Cas9 mediated knock-in in hPSCs

For Cas9-mediated reporter knock-in, ∼14,000 cells/cm^2^ hPSCs were seeded onto iMatrix-511 coated wells of a 12-well plate and cultured for 24 hours at 37°C, 5% CO_2_. For modRNA-based knock-in, iCas9 modRNA, target specific KI sgRNA, and the mini-donor plasmid were combined with Lipofectamine Stem Transfection Reagent (ThermoFisher) (1:4 ratio, mass/volume) in Opti-MEM medium (ThermoFisher). For plasmid-based knock-in, iCas9 modRNA was replaced with the EFS-Cas9 plasmid. Before transfection, the spent culture medium was replaced with fresh TeSR supplemented with 10 μM Y-27632. The transfection mix was incubated at room temperature (RT) for 10 minutes before being added to the well in a dropwise fashion. The transfection media was changed 24 hours later, and cells were maintained in TeSR with daily media changes.

### ModSAM-mediated target gene activation in hPSCs

For modSAM-mediated gene activation, ∼28,000 cells/cm^2^ hPSCs were seeded onto iMatrix-511 coated wells of a 12-well plate and cultured for 24 hours at 37°C, 5% CO_2_. For modRNA-based activation, the dCas9 activator (VP64dCas9VP64-2A-MPH) modRNA and target specific dgRNA were combined with Lipofectamine Stem Transfection Reagent (1:4 ratio, mass/volume) in Opti-MEM medium. For plasmid-based activation, the dCas9 activator modRNA was replaced with pPB-R1R2-EF1a-VP64dCas9VP64-T2A-MS2p65HSF1-IRES-bsd-pA plasmid (Addgene, #113341). Before transfection, the spent culture medium was replaced with fresh TeSR supplemented with 10 μM Y-27632. The transfection mix was incubated at RT for 10 minutes before being added to the well in a dropwise fashion followed by a media change 24 hours later. Cells were collected between 36 and 48 hours after transfection for either fluorescence activated cell-sorting (FACS) or flow cytometry.

### Fluorescence-activated cell-sorting (FACS)

Cells were washed with DPBS and dissociated into single cells with Accutase at 37°C for 10 minutes. Dissociated cells were transferred to excess DMEM (1:2, volume/volume) and spun down at 200x rcf for 5 minutes. After centrifugation, cells were resuspended in TeSR supplemented with 2% (volume/volume) Pen-Strep and 10 μM Y-27632. FACS was performed in either the Beckman Coulter MoFlo Astrios or Invitrogen Bigfoot Spectral Cell Sorter. Gated populations were collected in TeSR supplemented with 2% (volume/volume) Pen-Strep and 10 μM Y-27632. Cells were replated onto Matrigel (Corning) coated wells of a 48-well plate in TeSR supplemented with 2% (volume/volume) Pen-Strep, 10 μM Y-27632, and 0.75 μg/mL iMatrix-511.

### Flow Cytometry

Cells were washed with DPBS and dissociated into single cells with Accutase at 37°C for 10 minutes. Dissociated cells were transferred to excess DMEM (1:2, volume/volume) and spun down at 200x rcf for 5 minutes. After centrifugation, cells were resuspended in either Flow Buffer-1 (5% FBS in PBS) for live-cell analysis or fixed with 1% PFA in PBS at RT for 30 minutes and resuspended in Flow Buffer-2 (5% FBS + 0.1% Triton X-100 in PBS). If required, immunostaining was performed sequentially with appropriate primary and secondary antibodies (**Supplementary Table 9 and 10**). Data was collected on the BD Accuri C6 Plus flow cytometer and processed using the FlowJo software.

### Derivation of single-cell clones

Following FACS, sorted reporter hPSCs were seeded onto iMatrix-511 coated wells of a 6-well plate at ∼26 cells/cm^2^. Once the single-cell derived colonies had become sufficiently large, the wells were washed with DPBS and cells were incubated with DPBS for 2 minutes at RT. Single-cell derived colonies were picked manually with a 2-20 μl micro pipettor under the microscope and replated into individual iMatrix-511 coated wells of a 48-well plate in TeSR supplemented with 2% (volume/volume) Pen-Strep and 10 μM Y-27632.

### PCR genotyping of single-cell derived clones

Genomic DNA was extracted from single-cell derived clones with the Quick-DNA Miniprep Plus Kit (Zymo Research). The DNA concentration was measured using a NanoDrop. PCR reactions were set up with 100-200 ng of template DNA, 0.5 μM forward and reverse primers (**Table S7**), and Q5 High-Fidelity 2X Master Mix (New England Biolabs). 20 μL of PCR product was loaded and run on a 1% (weight/volume) agarose gel and imaged with the ChemiDoc Touch Imaging System (Bio-Rad).

### qPCR

hPSCs were lysed with TRI-Reagent and RNA was extracted using the Direct-zol RNA MiniPrep Plus kit (Zymo Research). RNA concentration was measured using a NanoDrop. cDNA was synthesized from extracted RNA using ZymoScript RT PreMix kit (Zymo Research). 2 ng of the cDNA template was combined with 0.25 μM forward and reverse qPCR primers (**Table S8**), and SYBR Green PCR master mix (ThermoFisher). The qPCR was run on a CFX Connect real-time qPCR machine (Bio-Rad). GAPDH was used as the reference gene and data was analyzed with ΔΔC_T_ method.

### Pancreatic Progenitor Differentiation

hPSCs were seeded onto iMatrix-511 coated wells and cultured in TeSR until cells were 90% confluent. Once the cells had reached the desired confluency, hPSCs were treated with optimized concentration of CHIR99021 in RPMI for 24 h (day 0 to day 1). From Day 1 to Day 4, cells were cultured in RPMI containing 2% v/v B-27 without insulin supplement (Thermo Fisher) with daily media changes. From day 4 to day 6 cells were cultured in RPMI supplemented with 0.05% HSA, 200 μg/ml ascorbic acid (Vc), 16 ug/ml Selenium, and 5 ug/ml Transferrin, with daily media changes. From day 6 to day 10, the cells were treated with 0.25 μM SAINT1, 2 μM RA, 0.75 μM Dorsomorphin and 0.2 μM PDBU in DMEM supplemented with 1% (volume/volume) B-27 and 50 μg/mL Vc; with media changes every other day. From day 10 to day 14 (Stage 4), cells were treated with 100 μM Y-27632, 10 mM nicotinamide, 0.75 μM Dorsomorphin and 100 ng/ml EGF in DMEM supplemented with 1% B-27 and 50 μg/ml Vc.

### Cardiac Differentiation

hPSCs were seeded onto Matrigel coated wells at 140,000 – 280,000 cells/cm^2^ and cultured in TeSR for 2-3 days. Cells were treated with an optimized concentration of CHIR99021 in RPMI supplemented with B-27 without insulin for 24 hours (day 0 to day 1). From day 1 to day 3 the cells were maintained in RPMI supplemented with B-27 without insulin. On day 3, cells were treated with 2 μM C59 in a 1:1 mixture of day 3 conditioned media and fresh RPMI supplemented with B-27 without insulin for 48 hours. From day 5 to day 7, cells were maintained in RPMI supplemented with B-27 without insulin. From day 7 onwards, cells were maintained in RPMI supplemented with B-27.

### Immunofluorescent Staining

Cells were washed with DPBS and fixed with 4% PFA in PBS at RT for 15 minutes. Cells were washed with PBS and blocked using 5% non-fat dry milk and 0.1% Triton X-100 in PBS for 1 hour. Following blocking, cells were sequentially stained using the appropriate primary and secondary antibodies (**Table S9, S10**). Hoechst 33342 (Thermofisher) was used to stain cell nuclei. Images were captured using a Nikon Ti Eclipse epifluorescence microscope and processed using the ImageJ software.

### Statistical Analysis

Quantification of flow cytometry data is shown as the mean ± SE unless otherwise specified. All statistical analyses were performed in R. Unpaired Student’s t-test was used for comparison between two groups. For comparison between multiple groups a two-way ANOVA followed by post-hoc Tukey’s test was used. A value of p > 0.05 was considered not significant; p < 0.05 (*), p < 0.01 (**), p < 0.001 (***) were considered statistically significant.

## Acknowledgments

This work was supported by NSF CBET-1943696 (to X.L.L.), NSF CBET-2143064 (to X.B.), NIH R21EB026035 (to X.L.L.), NIH R56DK133147 (to X.L.L.), and NIH R37CA265926 (to X.B.).

## Author contributions

T.H. and X.L.L. designed the experiments and analyzed the results. T.H. and J.L. performed the experiments and analyzed data. X.L.L. supervised the experiments. T.H. and X.L.L. wrote the manuscript. T.H., X.B., and X.L.L. contributed to the revision of the manuscript.

## Declaration of interests

All authors declare no competing interests.

**Figure S1 Transient activation of target gene can be used to purify fluorescent reporters.** (A-E) WT H9 cells were transfected with dCas9 activator modRNA and either SOX17 dgRNA or NKX6.1 dgRNA on day 0. On day 5 cells were stained for canonical stem cell markers OCT4 (A to E), NANOG (A), SOX2 (A), and TRA-1-60 (B to E) and analyzed by either immunofluorescent imaging (A) or flow cytometry (B to E). (A) Scale bar, 100 μm. (B) Representative flow cytometry plots showing OCT4 and TRA-1-60 expression untransfected WT H9 cells. (C) Representative flow cytometry plots showing expression of TRA-1-60 in day 5 post-transfection cells. (D) Representative flow cytometry plots showing expression of OCT4 in day 5 post-transfection cells. (E) Quantification of (C and D) with error bars representing the SEM across three independent replicates. (F) WT H9 cells were transfected with CD90 knockout sgRNA and either Cas9-2A-Puro modRNA or Cas9-2A-p53DD (iCas9) modRNA on day 0. Day 5 cells were collected and CD90 expression was analyzed by flow cytometry. Error bars represent the SEM of six independent replicates. p < 0.05 (*). (G-H) PDX1-EGFP reporter cells were generated from WT H9 cells as shown in (Figure 1 H). Cells were analyzed for EGFP expression by flow cytometry, 48 hours after activating PDX1 by either plasmid-based or modRNA-based approaches. (G) Representative flow cytometry plots showing EGFP expression across each of the four combinations. (H) Quantification of (G) with error bars representing SEM across 3 independent replicates. p < 0.05 (*), p < 0.01 (**), p < 0.001 (***).

**Figure S2 MAGIK for generating IMR90 PDX1-EGFP reporter cells.** (A) WT IMR90 iPSCs were transfected with iCas9 modRNA, PDX1 KI sgRNA, and PDX1-EGFP mini-donor plasmid. Cells were expanded in TeSR for several days before being passaged into multiple wells of a 12-well plate. The mixed knock-in cells were transfected with dCas9 activator modRNA and PDX1 dgRNA. EGFP^+^ cells were sorted via FACS 48 hours after transfection and replated into a single well of a 48-well plate. Representative plot showing percentage of EGFP positive cells sorted from total population. (B) Representative brightfield and fluorescent images of untransfected knock-in reporter cells. Scale bar, 100 μm. (C) Representative brightfield and fluorescent images of knock-in reporter cells before and after sorting until day 4 post-transfection. Scale bar, 100 μm. (D) Post-sort IMR90 PDX1-EGFP reporter cells were transfected with dCas9 activator modRNA and PDX1 dgRNA. On day 2 following transfection, cells were fixed and stained for PDX1 (Red). Representative fluorescent images showing co-localization of anti-PDX1 signal and EGFP. Scale bar, 100 μm. (E-F) Post-sort IMR90 PDX1-EGFP reporter cells were stained for pluripotency markers OCT4 (E and F), NANOG (E), SOX2 (E) and TRA-1-60 (F) and analyzed by either immunofluorescent imaging (E) or flow cytometry (F). (E) Scale bar, 100 μm. (F) Representative flow cytometry plots and quantification of OCT4 and TRA-1-60 expression in post-sort IMR90 PDX1-EGFP reporter cells. Error bars represent SEM across three replicates. (G) PCR genotyping of twelve single-cell derived clones from post-sort IMR90 PDX1-EGFP reporter cells. The expected bands from each set of probes are highlighted by a pair of red dashed lines (5’ OUT: 1746 bp; 3’ OUT: 1837 bp). (H) PCR genotyping of single-cell derived clones using Inner primers to distinguish between monoallelic and biallelic knock in (WT: 885 bp; KI with GFP: 1659 bp)

**Figure S3 MAGIK for generating H1 PDX1-EGFP reporter cells.** (A) WT H1 cells were transfected with iCas9 modRNA, PDX1 KI sgRNA, and PDX1-EGFP mini-donor plasmid. Cells were expanded in TeSR for several days before being passaged into multiple wells of a 12-well plate. The mixed knock-in cells were transfected with dCas9 activator modRNA and PDX1 dgRNA. EGFP^+^ cells were sorted via FACS 48 hours after transfection and replated into a single well of a 48-well plate. Representative plot showing percentage of EGFP positive cells sorted from total population. (B) Representative brightfield and fluorescent images of untransfected knock-in reporter cells. Scale bar, 100 μm. (C) Representative brightfield and fluorescent images of knock-in reporter cells before and after sorting until day 4 post-transfection. Scale bar, 100 μm. (D) Post-sort H1 PDX1-EGFP reporter cells were transfected with dCas9 activator modRNA and PDX1 dgRNA. On day 2 following transfection, cells were fixed and stained for PDX1 (Red). Representative fluorescent images showing co-localization of anti-PDX1 signal and EGFP. Scale bar, 100 μm. (E-F) Post-sort H1 PDX1-EGFP reporter cells were stained for pluripotency markers OCT4 (E and F), NANOG (E), SOX2 (E) and TRA-1-60 (F) and analyzed by either immunofluorescent imaging (E) or flow cytometry (F). (E) Scale bar, 100 μm. (F) Representative flow cytometry plots and quantification of OCT4 and TRA-1-60 expression in post-sort H1 PDX1-EGFP reporter cells. Error bars represent SEM across three replicates. (G) PCR genotyping of single-cell derived clones from post-sort H1 PDX1-EGFP reporter cells. The expected bands from each set of probes are highlighted by a pair of red dashed lines (5’ OUT: 1746 bp; 3’ OUT: 1837 bp). (H) PCR genotyping of single-cell derived clones using Inner primers to distinguish between monoallelic and biallelic knock in (WT: 885 bp; KI with GFP: 1659 bp)

**Figure S4 MAGIK for generating 6-9-9 PDX1-EGFP reporter cells.** (A) WT 6-9-9 cells were transfected with iCas9 modRNA, PDX1 KI sgRNA, and PDX1-EGFP mini-donor plasmid. Cells were expanded in TeSR for several days before being passaged into multiple wells of a 12-well plate. The mixed knock-in cells were transfected with dCas9 activator modRNA and PDX1 dgRNA. EGFP^+^ cells were sorted via FACS 48 hours after transfection and replated into a single well of a 48-well plate. Representative plot showing percentage of EGFP positive cells sorted from total population. (B) Representative brightfield and fluorescent images of untransfected knock-in reporter cells. Scale bar, 100 μm. (C) Representative brightfield and fluorescent images of knock-in reporter cells before and after sorting until day 7 post-transfection. Scale bar, 100 μm. (D) Post-sort 6-9-9 PDX1-EGFP reporter cells were transfected with dCas9 activator modRNA and PDX1 dgRNA. On day 2 following transfection, cells were fixed and stained for PDX1 (Red). Representative fluorescent images showing co-localization of anti-PDX1 signal and GFP. Scale bar, 100 μm. (E-F) Post-sort 6-9-9 PDX1-EGFP reporter cells were stained for pluripotency markers OCT4 (E and F), NANOG (E), SOX2 (E) and TRA-1-60 (F) and analyzed by either immunofluorescent imaging (E) or flow cytometry (F). (E) Scale bar, 100 μm. (F) Representative flow cytometry plots and quantification of OCT4 and TRA-1-60 expression in post-sort 6-9-9 PDX1-EGFP reporter cells. Error bars represent SEM across three replicates.

