## Supplementary figures and images for "MAGIK: A rapid and efficient method to create lineage-specific reporters in human pluripotent stem cells"

### Fig S1

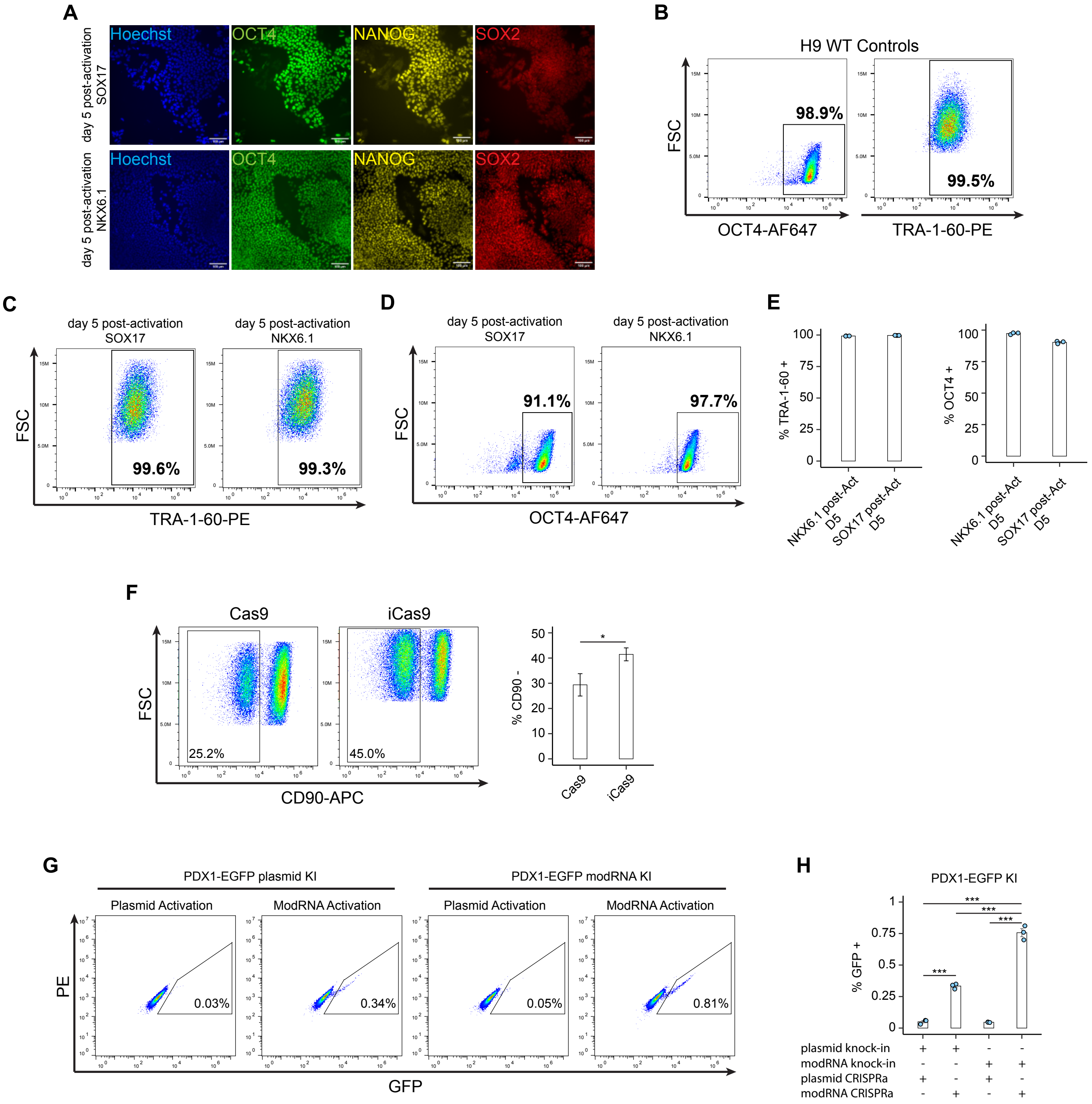

### Fig S2

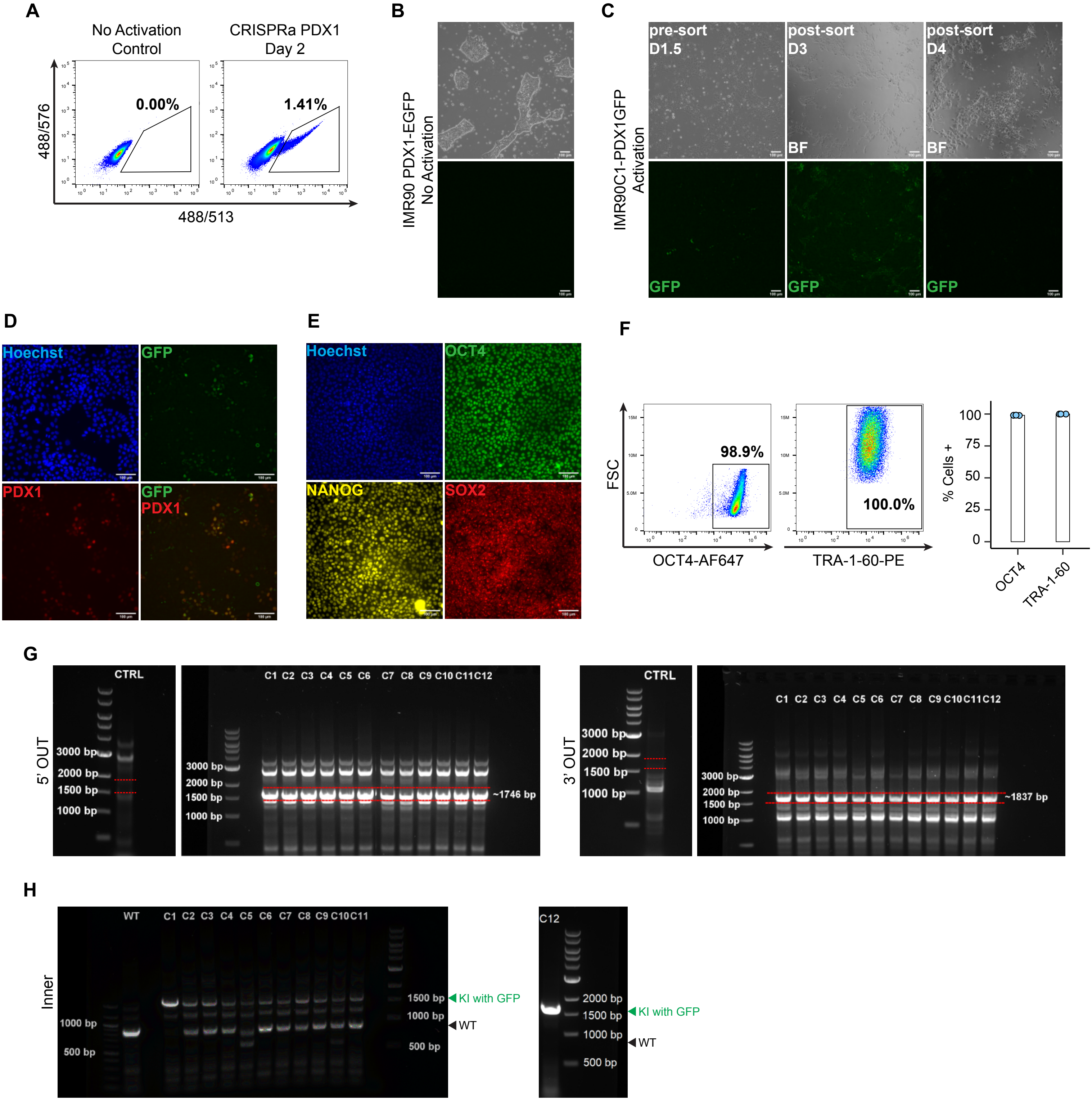

### Fig S3

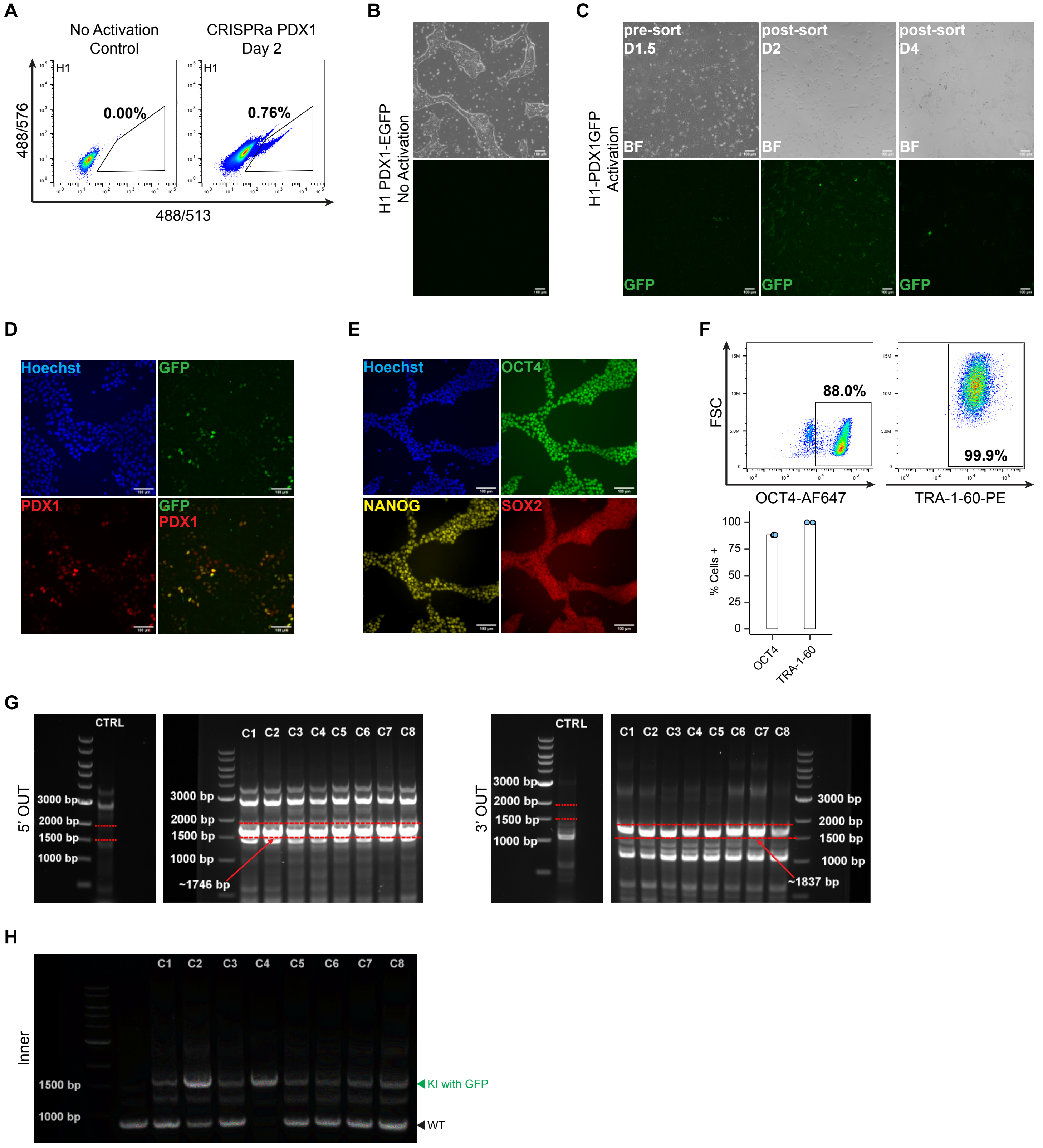

### Fig S4

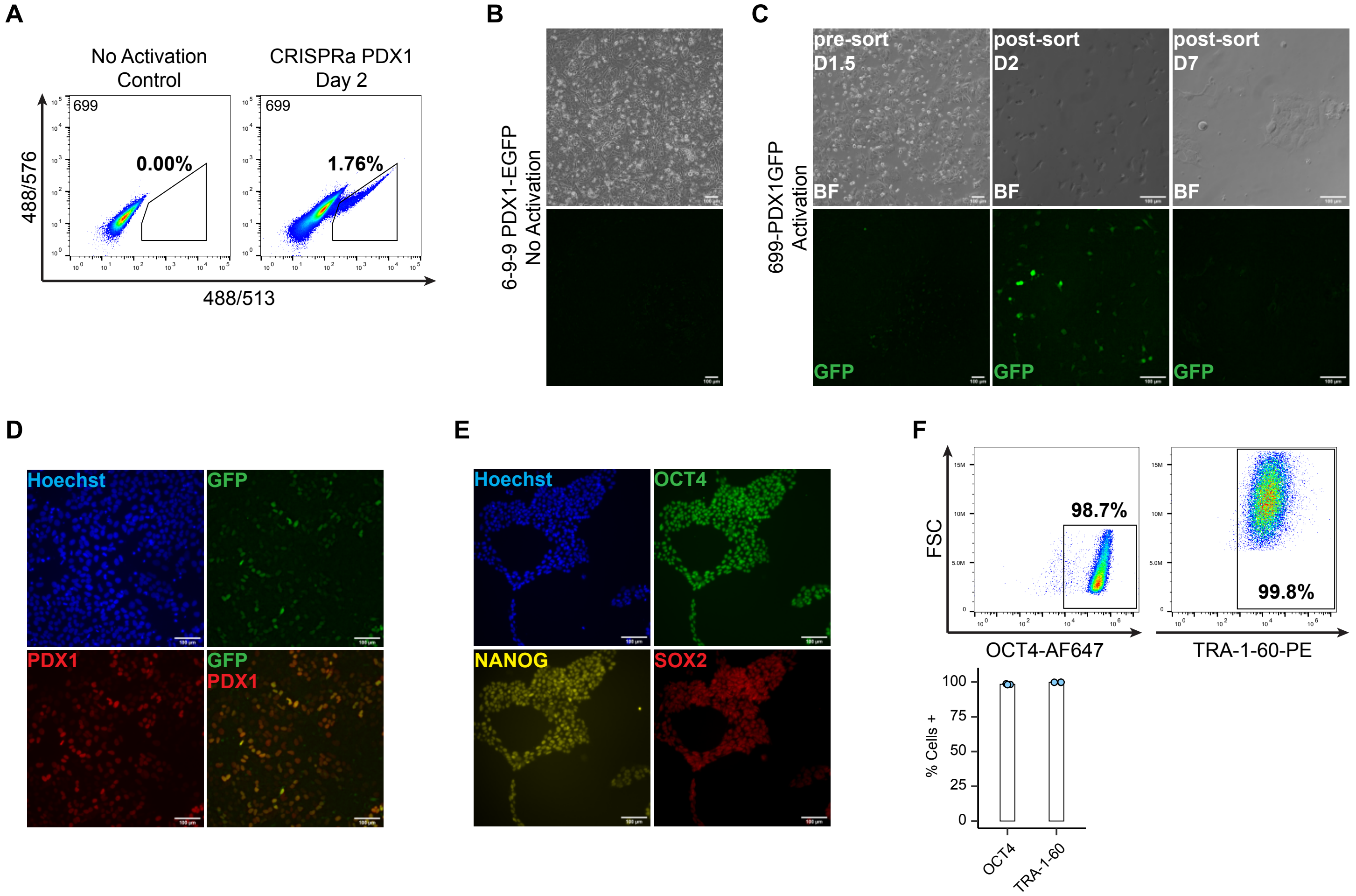
